# Rapid plant-to-plant systemic signaling via a *Cuscuta* bridge

**DOI:** 10.1101/2023.07.13.548730

**Authors:** Yosef Fichman, María Ángeles Peláez-Vico, Asha Kaluwella Mudalige, Hyun-Oh Lee, Ron Mittler, So-Yon Park

## Abstract

*Cuscuta*, commonly known as dodder, is a parasitic plant that thrives by attaching itself to the stems of other plants. It is found across the globe and is notorious for obstructing crop growth as a weed. Over the past decade, *Cuscuta* has been used to gain insights into plant-plant interactions and molecular trafficking. Here, we report that two plants connected via a *Cuscuta* bridge can exchange rapid systemic calcium, electric, and reactive oxygen species signals. These findings suggest that plant interactions with Cuscuta may have beneficial effects to plants, enabling them to rapidly communicate with each other.

## Manuscript

The ability of plants to rapidly transmit signals from one tissue (*e*.*g*., a leaf) to another (*e*.*g*., another leaf, roots, and/or reproductive tissues), termed ‘systemic signaling’, plays a key role in optimizing the plant overall photosynthetic activity, growth, productivity, and responses to abiotic and biotic stresses (Kollist et al. 2019). Among the different signals that mediate rapid systemic intra-plant tissue-to-tissue communication are electric, calcium, reactive oxygen species (ROS), and hydraulic waves (*e*.*g*., Miller et al. 2009; Mousavi et al. 2013; Toyota et al. 2018; Grenzi et al. 2023). These travel at rates of 0.5-to-several cm per minute, mostly via the plant vascular system, and carry information that triggers transcriptomic, metabolomic, proteomic, and physiological responses in systemic tissues (*e*.*g*., Suzuki et al. 2013; Nguyen et al. 2018; Zandalinas et al. 2020).

In recent years, it was found that systemic signals can also travel from plant-to-plant (inter-plant), below or above ground, and convey important information that coordinates the response of different plants living in a community to stress (*e*.*g*., Venkateshwaran et al. 2013; Szechyńska-Hebda et al. 2022). However, the potential of parasitic plants, that are generally considered to be pests, to mediate rapid plant-to-plant communication in response to stress is largely unknown. Parasitic plants such as dodder (*Cuscuta campestris*) are thought to ‘steal’ water and nutrients from host plants after forming physical connections with them, called haustoria, without providing any benefits back to the plant (Hibberd & Dieter Jeschke, 2001). Over the past decade, different studies revealed the transfer of mRNAs, small RNAs, DNA, and proteins between the host and *Cuscuta* (Jhu & Sinha, 2022). Mobility of mRNAs and proteins between two different plants connected by *Cuscuta* has also been shown (Liu et al. 2020), as well as the transfer of macromolecules associated with systemic herbivory (Hettenhausen et al. 2017; Zhuang et al. 2018). However, the potential of *Cuscuta* to transmit important rapid systemic signals, such as calcium, ROS, and membrane potential depolarization waves, between different plants in response to abiotic stress remains uncertain, prompting us to investigate the transmission of such inter-plant signals between two different host plants connected via a *Cuscuta* (that functions as a ‘bridge’).

To establish an experimental system to study the transfer of rapid systemic signals between two different plants connected by a *Cuscuta* bridge, we produced an Arabidopsis (*Arabidopsis thaliana*) ‘donor’ - *Cuscuta* ‘bridge’ – and ‘receiver’ *Arabidopsis*, inter-plant (plant-*Cuscuta*-plant) system (Fig. 1A) and studied the propagation of the ROS wave in this system. Both donor and receiver *Arabidopsis* were connected to *Cuscuta* through fully developed haustoria (Fig. 1B, black arrow). Using a live whole-plant imaging system (IVIS Lumina S5; Fichman et al. 2019), we then tracked the spread of the ROS wave within the inter-plant system after wounding a single rosette leaf of the donor *Arabidopsis* plant (Fig. 1A, yellow lightning bolt; This treatment was previously shown to send a systemic signal to neighboring leaves connected via the vascular system, and to the entire plant; Fichman et al, 2019). Interestingly, the accumulation of the systemic ROS signal was not limited to the local leaf of the donor plant but rather spread to and throughout the donor systemic leaves, *Cuscuta* bridge, and the entire receiver plant (Fig. 1C). This finding suggested that wounding of a single leaf of the donor plant triggered a systemic ROS wave that traveled through the *Cuscuta* bridge and triggered ROS accumulation in the receiver plant.

**Figure 1.**
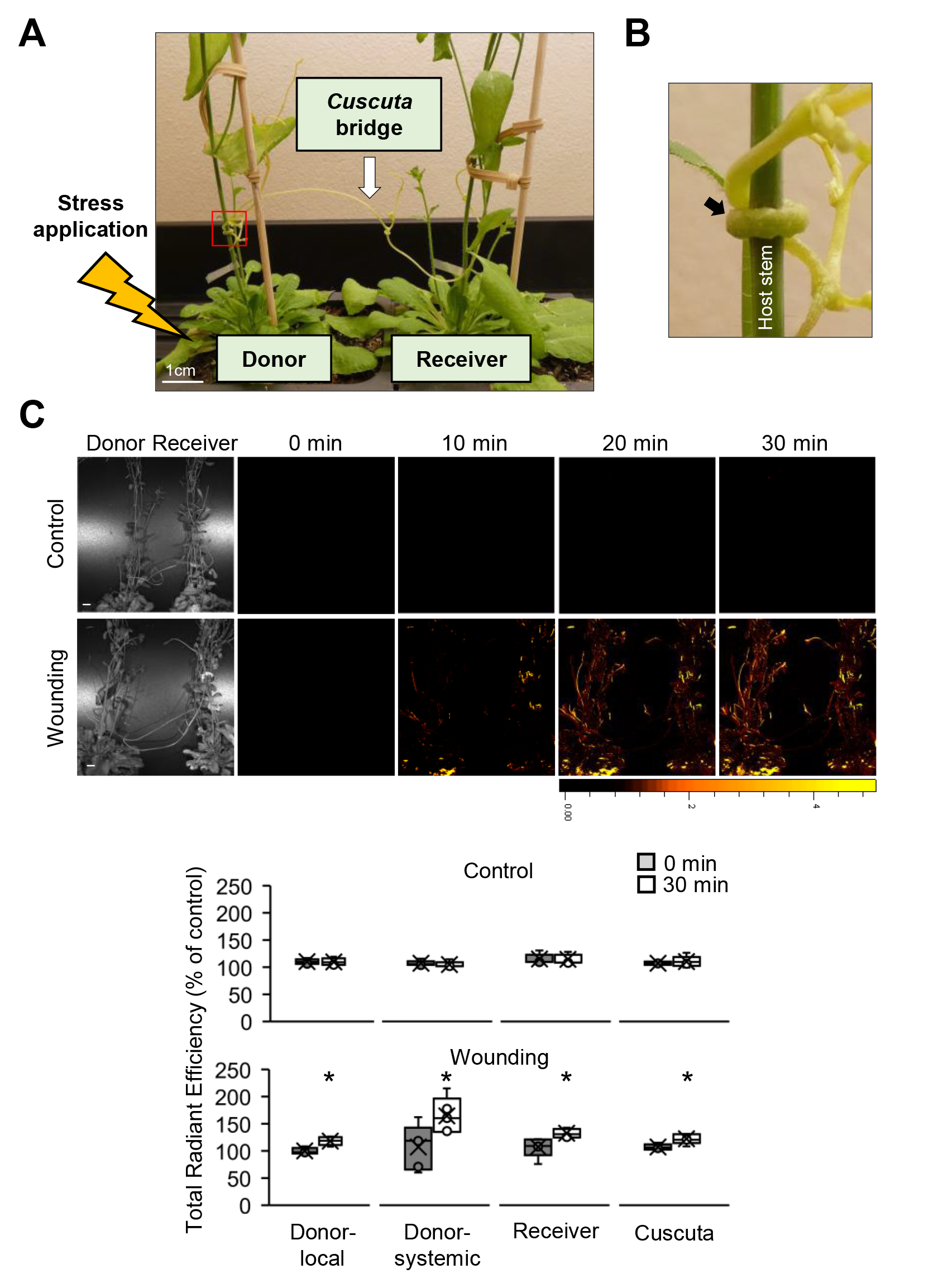
The experimental design used for measuring ROS accumulation following wounding. **A**. A *Cuscuta* bridge (white arrow) connected two *Arabidopsis* plants (donor *Arabidopsis* on the left and receiver *Arabidopsis* on the right); stress was applied to a single leaf of the donor plant (jagged yellow arrow). **B**. Enlarged image of the red box in A. The black arrow indicates established haustorium developed on the *Arabidopsis* stem. **C**. Representative 30 min time lapse images of ROS accumulation in control plants or following mechanical wounding stress applied to a single leaf of the donor plant (Top), and Quantification of fluorescence corresponding to ROS content at 0 min (grey) and after 30 min (white) in control treatment and following an injury of a single leaf of the donor plants, in donor’s local and systemic leaves, and in the receiver plant and *Cuscuta* bridge (Bottom). Asterisks indicated significance; Student’s t test (N=5; *P < 0.05). Results are displayed as box-and-whisker plots, with the borders corresponding to the 25th and 75th percentiles of the data. Each data value is included as a point within each box plot, with the horizontal line representing the median and “X” corresponding to the mean. Whiskers represent 1.5 times the minimum and maximum of the mean (1.5 times of the interquartile range); the size bar indicates 1 cm and is applicable to all other fluorescence images; units of color scale bar are total counts of fluorescence; fluorescence units used to calculate % of control are (p/s]/[µW/cm^2^; Fichman & Mittler, 2021).

Using the system developed in Fig. 1, we next examined the propagation of other rapid systemic signals from plant-to-plant through the *Cuscuta* bridge. To prevent potential interference from ROS signaling processes associated with the penetration of the *Cuscuta* haustoria into the host stems (which is a type of wounding stress; Johnsen et al. 2015; Hegenauer et al. 2016; Slaby et al. 2021), we used high light stress as a trigger of the different rapid systemic waves (Fig. 2). Applying high light stress to a single rosette leaf of a donor *Arabidopsis* plant led to ROS accumulation within 30 min in the entire plant-*Cuscuta*-plant chain including the donor, *Cuscuta*, and receiver plants (Fig. 2A; Video 1; representative of 5 different experiments; in all videos, color scale shows increase and decrease of the signal; zero is black; Fichman et al., 2021). In contrast, when *Cuscuta*-inoculated donor and *Cuscuta*-inoculated receiver plants were not connected via a *Cuscuta* bridge, the ROS signal did not cross over from the donor to the receiver plant, indicating that, at least under the conditions tested, volatile signals were not involved in the transmission of the plant-to-plant signal (Supplemental Fig. S1; for this analysis we also directly compared the intensity of signals between receiver plants in Fig. 2A and S1A; Fig S1C). As the ROS wave is frequently accompanied by the propagation of systemic calcium and electric waves, we repeated our experiments using the same plant-*Cuscuta*-plant chain system, however, using dye indicators for calcium and membrane potential (electric) waves (Fichman & Mittler, 2021; Fichman et al. 2022). Similar to the ROS wave observation (Fig. 2A; Video 1), we were able to image and measure the propagation of a calcium wave (Fig. 2B; Video 2; representative of 5 different experiments), and a membrane depolarization wave (Fig. 2C; Video 3; representative of 5 different experiments), across the entire plant-*Cuscuta*-plant system, in response to the local application of high light stress to a single leaf of the donor plant. Please note that soil bacteria/algae, as well as dead leaves can produce a fluorescence signal that would be visible in videos 1-3. These are not associated with the plant-to-plant signal and should be considered as background signals. Taken together, our findings reveal that ROS, calcium, and membrane potential waves, activated by a high light stress treatment applied to a single tissue (leaf) of the donor plant, are transmitted to a receiver plant through the *Cuscuta* bridge. These results indicate that systemic stress signals (ROS, calcium, and electric waves) can be transferred within minutes aboveground between different plants connected through a parasitic plant such as *Cuscuta*.

**Figure 2.**
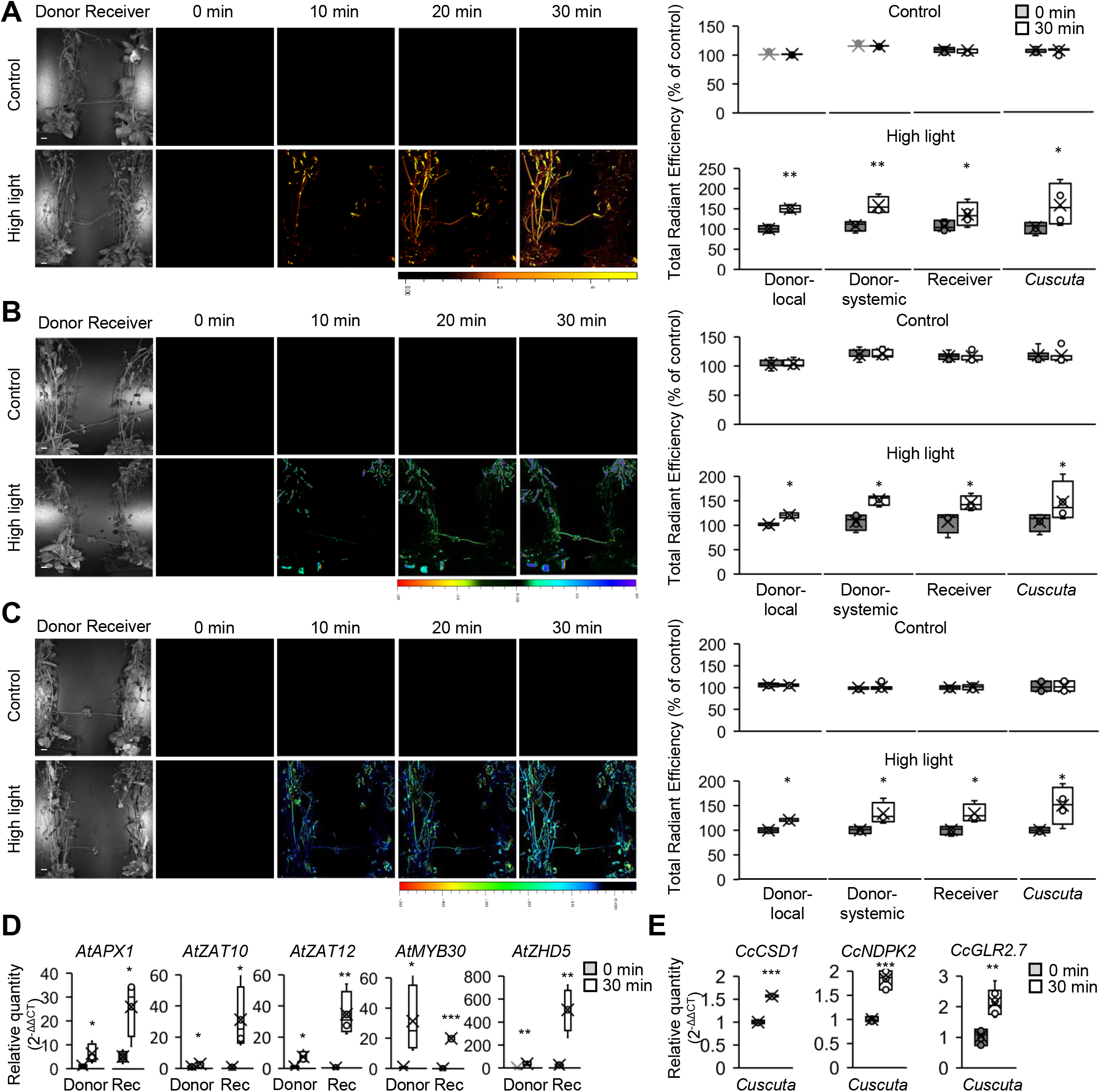
ROS accumulation and transcriptional changes in donor plant, *Cuscuta* bridge, and receiver plant, in response to a local light stress applied to a single leaf of the donor plant. **A**. Representative images of ROS accumulation in control and high light stress-treated donor plant, in a donor-*Cuscuta* bridge-receiver chain (light stress was applied to a single leaf of the donor plant; Left), and quantification of fluorescence corresponding to ROS content at 0 min (grey) and after 30 min (white) in control and following high light treatment applied to a single leaf of the donor plants (local leaf), in *Arabidopsis* donor’s local and systemic leaves, receiver and *Cuscuta* plants (Right). **B**. Similar to A, but for Ca^2+^ accumulation. **C**. Similar to A, but for membrane potential changes. **D**. Changes in the steady-state level of transcripts associated with ROS responses in *Arabidopsis* donor and receiver (Rec) plants, measured under control conditions (0 min; grey) or following treatment (30 min after light stress to a single leaf of the donor plant; white). **E**. Same as D, but for *Cuscuta* ROS-response transcripts. Results are shown as box-and-whisker plots, with the borders corresponding to the 25th and 75th percentiles of the data. Each data value is included as a point within each box plot, with the horizontal line representing the median and “X” corresponding to the mean. Whiskers represent 1.5 times the minimum and maximum of the mean (1.5 times of the interquartile range). Quantitative real-time results are presented as relative quantity PCR, normalized to reference gene. N=5-12, Asterisks indicated significant difference; Student’s t-test (N=7, 5, and 5, in A, B, and C, respectively; *P < 0.05, **P < 0.01 and ***P<0.001); the size bar indicates 1 cm and is applicable to other fluorescence images; units of color scale are total counts of fluorescence; fluorescence units used to calculate % of control are (p/s]/[µW/cm^2^; Fichman & Mittler, 2021). Abbreviations: *APX1, ASCORBATE PEROXIDASE 1; CSD1, COPPER/ZINC SUPEROXIDE DISMUTASE 1*; *GLR2*.*7, GLUTAMATE RECEPTOR 2*.*7; MYB30, MYELOBLASTOSIS DOMAIN PROTEIN 30; NDPK2, NUCLEOSIDE DIPHOSPHATE KINASE 2;* Rec, Receiver; *ZAT10, ZINC FINGER OF ARABIDOPSIS THALIANA 10; ZAT12, ZINC FINGER OF ARABIDOPSIS THALIANA 12; ZHD5, ZINC FINGER HOMEODOMAIN 5*.

To investigate whether the transmission of the different waves is associated with the triggering of molecular responses in the receiver plant, we performed RT-qPCRs using total RNA extracted from the different *Arabidopsis* and *Cuscuta* plants (Fig. 1A). Tissues were harvested before and after the application of the high light stress to the local tissue, and transcript levels were compared relative to the control condition. The qRT-PCRs results revealed an increase in the expression of *ASCORBATE PEROXIDASE 1* (*AtAPX1*), *MYELOBLASTOSIS DOMAIN PROTEIN 30* (*AtMYB30*), *ZINC FINGER OF ARABIDOPSIS THALIANA 10* (*AtZAT10*), and *ZINC FINGER HOMEODOMAIN 5* (*AtZHD5*) transcripts in both donor and receiver *Arabidopsis* plants under the high light stress (Fig. 2D). These transcripts are known to accumulate in local and systemic tissues of *Arabidopsis* in response to a 2 min local light stress (Fichman et al. 2022). The enhanced transcript expression in receiver plants suggest that the propagation of systemic signals through the *Cuscuta* bridge stimulates a molecular response in them. Furthermore, we investigated changes in the steady-state levels of the stress-associated *Cuscuta* transcripts, *COPPER/ZINC SUPEROXIDE DISMUTASE 1* (*CcCSD1*), *NUCLEOSIDE DIPHOSPHATE KINASE 2* (*CcNDPK2*), and *GLUTAMATE RECEPTOR 2*.7 (*CcGLR2*.*7*), to understand the potential influence of the systemic waves on *Cuscuta*. Our qRT-PCR result revealed significant changes in the expression of these transcripts (Fig. 2E), indicating that the transmitted systemic signals from the host plant could also trigger transcriptional responses in *Cuscuta*.

When viewing the data shown in Fig. 1C and 2A-2C, it is important to note that different plant tissues can absorb the different imaging dyes, and/or respond to/metabolize ROS, calcium, and/or membrane potential at different rates, resulting in differential intensities and rates of signal imaging/propagation. For example, seed pods might absorb the dye much faster, and/or display much higher rates of ROS production/much lower rates of ROS scavenging. In addition, as we previously demonstrated (Fichman et al. 2019), mutants deficient in ASCORBATE PEROXIDASE 1, that scavenges ROS, develop the ROS signal much faster than wild type, and mutants deficient in RESPIRATORY BURST OXIDASE HOMOLOG (RBOH) D/F develop the ROS signal much slower, as they produce less signaling ROS. The expression level of certain genes in specific tissues, may therefore further influence the detection of the ROS signal. As we know very little about *Cuscuta* ROS metabolism, compared to *Arabidopsis*, the rates of signal detection may also be slower in *Cuscuta* due to similar reasons (*i*.*e*., dye uptake, ROS scavenging/production/transport, etc). As we demonstrate the plant-to-plant signal transmission via 3 different dyes, with control experiments, and using qRT-PCR analyses (Fig. 1, 2, and Supplemental Fig. S1), we are confident that our findings reveal the transmission of different plant-to-plant signals via a *Cuscuta* bridge, and that the different imaging intensities are simply a result of differences in dyes uptake and/or rates of ROS/calcium/membrane potential metabolism/signaling between the different tissues. In future studies, we would like to test the transfer of the plant-to-plant signal via a *Cuscuta* bridge using different *Cuscuta* mutants deficient in RBOHs or different ROS metabolism enzymes, as well as identify the source of ROS produced in *Cuscuta* and the two different plants during this response (other than RBOHs).

In a recent study, we demonstrated that under humid conditions plants growing in a community can transfer ROS and electric wave signals between each other, given that their leaves are physically touching each other (Szechyńska-Hebda et al. 2022). Here we extend our findings and report that individual plants that are connected through the parasitic plant *Cuscuta* can exchange systemic ROS, calcium, and membrane potential signals in response to local stresses such as wounding or high light stress. As the transmission of systemic stress signals between different plants connected by a *Cuscuta* bridge could activate acclimation mechanisms (Fig. 2D, 2E) and contribute to the survival of these plants (and/or the entire community) during stress, it is possible that plant interactions with *Cuscuta* may provide certain benefits to plants, especially plants growing within a community. This possibility should be addressed in future studies as it may change the definition of *Cuscuta*/dodder from a parasitic plant to a partially mutualistic symbiotic plant.

## Supplementary Data

**Supplemental Figure S1**. ROS accumulation in donor, but not receiver plant, when the two *Cuscuta*-infected plants are not connected via a *Cuscuta* bridge.

**Supplementary Table S1**. List of primers used in this study.

**Methods S1**. Planting materials and germination of *Arabidopsis*

**Methods S2**. Introduction of *Cuscuta* tissues

**Methods S3**. Facilitating parasitic connection between two *Arabidopsis* plants

**Methods S4**. Wounding and high light stress application

**Methods S5**. ROS, calcium, and membrane depolarization measurements

**Methods S6**. RNA extraction and qRT-PCRs

**Methods S7**. Statistical analysis

## Author contributions

SP, YF, and RM conceived the ideas and designed the experiments. YF, AM, MAPV, and HL conducted the experiments and analyzed the data. SP, YF, and RM wrote the manuscript. All authors contributed critically to the drafts and gave final approval for publication.

## Data availability

The datasets used and/or analyzed during the current study are available from the corresponding author upon request.

## Figure Legends

**Video 1**. ROS wave in response to high light stress

**Video 2**. Calcium wave in response to high light stress

**Video 3**. Membrane depolarization wave in response to high light stress

