## Supplementary data for "Rapid plant-to-plant systemic signaling via a *Cuscuta* bridge"

The following Supporting Information is available for this article:

**Supplementary Figure 1.** ROS accumulation in donor, but not receiver plant, when the two *Cuscuta*-infected plants are not connected via a *Cuscuta* bridge.

**Supplementary Table1.** List of primers used in this study.

**Methods S1**. Planting materials and germination of *Arabidopsis*

**Methods S2**. Introduction of *Cuscuta* tissues

**Methods S3**. Facilitating parasitic connection between two *Arabidopsis* plants

**Methods S4**. Stress application

**Methods S5**. ROS, Calcium, and membrane depolarization measurements

**Methods S6**. RNA extraction and qRT-PCRs

**Methods S7**. Statistical analysis

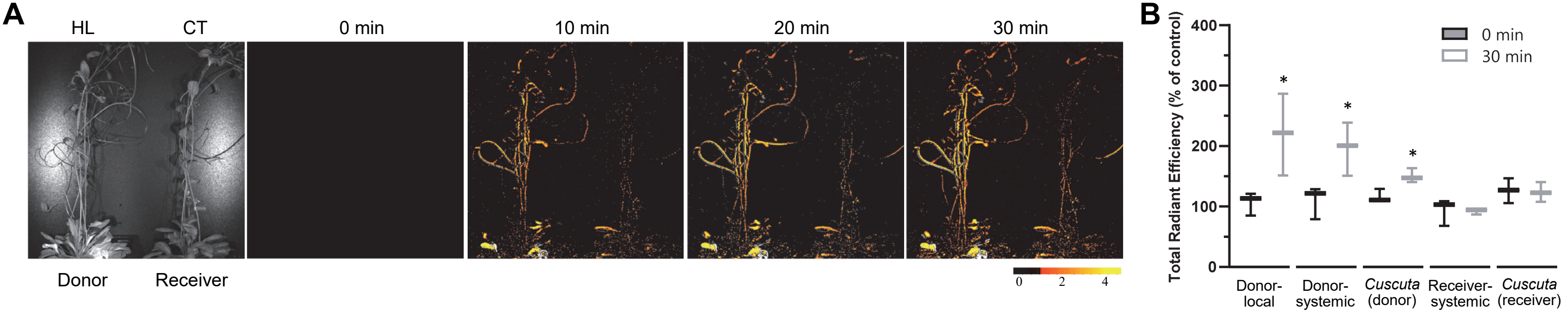


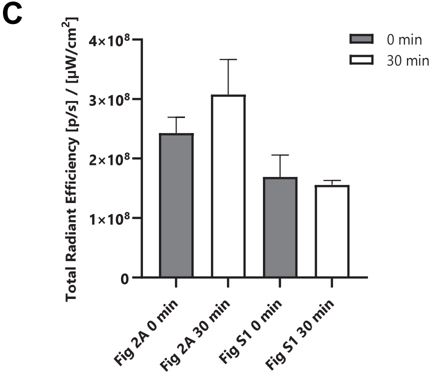


**Supplemental Figure S1.** ROS accumulation in donor, but not receiver plant, when the two Cuscuta-infected plants are not connected via a Cuscuta bridge. A. Representative images of ROS accumulation in a high light stress-treated Cuscuta-infected donor plant, but not a Cuscuta-infected receiver plant, when the two plants are not connected by a Cuscuta bridge, over 30 min. Abbreviations: CT, control; HL, high light; min, minutes. B. Quantification of fluorescence corresponding to ROS content at 0 min (black) and after 30 min (grey) in local and systemic leaves of donor and receiver, and in Cuscuta attached to donor or receiver plants that are not connected by a Cuscuta bridge. Box and whisker plots show 25th and 75th percentiles of 3 experiments. Center line corresponds to the median and the whiskers to the maximum and minimum values. Asterisks denote significant differences between treated and un-treated plants (N = 3; $*P < 0.05$, Student's t-test). Units of color scale are total counts of fluorescence; fluorescence units used to calculate % of control in B are (p/s]/[µW/cm2; Fichman & Mittler, 2021). C. As each image generated for each experiment is based on the overall total signal included in each individual experiment, to directly compare the signal intensity between the receiver plants in Figures 2A and S1, we used the raw intensity reads of each experiment. These values were not adjusted to the overall reads from each experiment and could therefore directly compare the signal in a receiver plant connected via a Cuscuta bridge to a stressed plant (Figure 2A) and a receiver plant not connected by a bridge to a stressed plant (Figure S1A). As shown in the bar graph presented in Figure S1C, the signal intensity of a receiver plant not connected by a Cuscuta bridge was not significantly different from that of a control untreated plant. The graph represents the means ± SE of three independent experiments.

**Supplementary Table S1.** List of primers used in this study.

| Plants | Note | Gene name (accession number) | Forward | Reserve |
|---|---|---|---|---|
| *Arabidopsis* | Target | *AtAPX2* (AT3G09640) | TCATCCTGGTAGACTGGACAAA | CACATCTCTTAGATGATCCACACC |
| | Target | *AtMYB30* (AT3G28910) | CCACTTGGCGAAAAAGGCTC | ACCCGCTAGCTGAGGAAGTA |
| | Target | *AtZAT10* (AT1G27730) | ACTAGCCACGTTAGCAGTAGC | GTTGAAGTTTGACCGGAAGTC |
| | Target | *AtZAT12* (AT5G59820) | TGGGAAGAGAGTGGCTTGTTT | TAAACTGTTCTTCCAAGCTCCA |
| | Target | *AtZHD5* (AT1G75240) | CCACCAATCCAAGTCTCCCTC | GCTCGCCGCATGATTCTTTAG |
| | Reference | *AtEF1a* (AT5G60390) | GAGCCCAAGTTTTTGAAGA | CTAACAGCGAAACGTCCCA |
| *Cuscuta* | Target | *CcNDPK2* (Cc000774) | GGTTTGAAGCTCATCACTGTCG | CACAACAGGCCCAGAAACAATG |
| | Target | *CcGLR2.7* (Cc025976) | CGTCTTTGCTCACAGGGAAAAG | TAACATGGACGTTAAGCTGGCT |
| | Target | *CcCSD1* (Cc004705) | GAATCATCATGGGGCTCCTGAT | GGTCCAGTGAGAGGAATCTGAC |
| | Reference | *CcTubulin* (Cc015968) | AGTTCCAGACCAATCTTGTCCC | TGAGACAGCAAGCCATGTACTT |

**Methods S1**. Planting materials and germination of *Arabidopsis*

*Arabidopsis* (Columbia-0) seeds were sterilized with 30% commercial bleach (NaOCl) and 0.02% Triton-X100 with autoclaved water. Sterile seeds were soaked in 1ml water in 4°C for 24 hours and placed on Gamborg B5 media. All plates containing *Arabidopsis* seeds were incubated in a chamber with 16 hours light / 8 hours dark with 55 µmol $m^{-2}$ $s^{-1}$ light intensity, 22-24°C conditions. Eight days after germination, the *Arabidopsis* seedlings were transferred to the soil pots (Promix BX) and maintained in the same conditions.

**Methods S2**. Introduction of *Cuscuta* tissues

For nursery *Cuscuta* stems, seedlings of the lab-grown line of *Cuscuta campestris* were grown on beets (Beta vulgaris) for one month in greenhouse conditions (30°C with 14 hours light and 10 hours dark). When the *Arabidopsis* plants started to form the main flowering stem, 3-4 cm long *Cuscuta* shoot tip with apical node were attached to the main flowering stem of donor *Arabidopsis* plant. After 24-48 hours, the *Cuscuta* coiled around the *Arabidopsis* stems. *Cuscuta* began haustoria formation 3-4 days after attachment and grew new shoots 7-8 days after attachment.

**Methods S3**. Facilitating parasitic connection between two *Arabidopsis* plants

When the new *Cuscuta* shoot growing on an initial *Arabidopsis* plant (donor plant) reached about 10 cm long, it was connected to a new *Arabidopsis* plant (receiver plant). The connection was facilitated by attaching the new *Cuscuta* shoot to the flowering stem of a receiver *Arabidopsis* plant, forming a *Cuscuta* connection bridge between the two hosts. After 4-5 days of attachment, the *Cuscuta* vine coiled around the new host stem, created haustoria, and formed new tissues, creating the inter-plant system.

**Methods S4**. Wounding and high light stress application

Wounding stress was induced by simultaneously puncturing a single leaf from the donor *Arabidopsis* with 18 dressmaker pins. Cool white light of 1200 µmol photons $s^{-1}$ $m^{-2}$ generated by fiber optic (Schott) was applied to a single donor plant leaf for 2 min. The waves and RNA sampling followed as described below.

**Methods S5**. ROS, calcium, and membrane depolarization measurements

The donor, receiver, and *Cuscuta* inter-plant system was fumigated with a solution of 0.1 M phosphate buffer Ph 7.4 containing $H_2$DCF-DA (for ROS measurement), Flou-4-AM (Sigma; for cytosolic calcium), or DiBAC3(4) (Biotium; for depolarization of the plasma membranes) and 0.001% Silwet L-77 (bioWORLD) for 30 min using nebulizers (Punasi Direct). Following the fumigation, high light stress was applied to a single leaf. The connected plants were then placed in the IVIS for 30 min for fluorescence detection (ex./em. 480 nm/ 520 nm). In the control treatment, the connected plants were put in the IVIS after the fumigation, without stress treatment. The results were analyzed with Living Image software, using the math function to subtract the threshold fluorescence.

**Methods S6**. RNA extraction and qRT-PCRs

Tissue samples from donor plant, receiver plant, and *Cuscuta* were collected at either 0 min or 30 min after local high light treatment to the donor plant. Collected tissues were ground and total RNA was extracted using the Plant RNeasy Kit (Qiagen). cDNAs (PrimeScript RT Reagent Kit; Takara Bio) were synthesized, and qRT-PCRs were performed using the Bio-Rad IQ mix and gene specific primers (Supplementary Table 1). Relative quantification of target genes was calculated using $2^{-\Delta\Delta CT}$ method after normalizing with reference genes and the control condition (0 min after local high light to donor plant).

**Methods S7**. Statistical analysis

All experiments were repeated at least three times. Box plot graphs were presented with the mean as X, median denoted as the line in the box, and box borders are 25th and 75th percentiles; whiskers are the 1.5 interquartile range. P-values ($^*P < 0.05$, $^{**}P < 0.01$, $^{***}P < 0.001$) were generated with two-tailed Student *t* test paired samples.
